# Incorporating cell hierarchy to decipher the functional diversity of single cells

**DOI:** 10.1101/2022.08.17.504240

**Authors:** Lingxi Chen, Shuai Cheng Li

## Abstract

Cells possess functional diversity hierarchically. However, most single-cell analyses neglect the nested structures while detecting and visualizing the functional diversity. Here, we incorporate cell hierarchy to study functional diversity at subpopulation, club (i.e., sub-subpopulation), and cell layers. Accordingly, we implement a package, SEAT, to construct cell hierarchies utilizing structure entropy by minimizing the global uncertainty in cell-cell graphs. With cell hierarchies, SEAT deciphers functional diversity in 36 datasets covering scRNA, scDNA, scATAC, and scRNA-scATAC multiome. First, SEAT finds optimal cell subpopulations with high clustering accuracy. It identifies cell types or fates from omics profiles and boosts accuracy from 0.34 to 1. Second, SEAT detects insightful functional diversity among cell clubs. The hierarchy of breast cancer cells reveals that the specific tumor cell club drives *AREG*-*EGFT* signaling. We identify a dense co-accessibility network of *cis*-regulatory elements specified by one cell club in GM12878. Third, the cell order from the hierarchy infers periodic pseudo-time of cells, improving accuracy from 0.79 to 0.89. Moreover, we incorporate cell hierarchy layers as prior knowledge to refine nonlinear dimension reduction, enabling us to visualize hierarchical cell layouts in low-dimensional space.

## Introduction

Cells in the biological system own functional diversity hierarchically, which signifies cell types or states during development, disease, and evolution, up to the biosystem (1, 2). The heterogeneity of the cell is observed with nested structures (3). In the tumor microenvironment, infiltrated lymphocytes include B cells and T cells. Furthermore, T cells can be classified into helper T cells and cytotoxic T cells (4). Specific expression of the marker genes *CD4* and *CD8* will strengthen intra-similarity within helper and cytotoxic T cells, respectively, resulting in nested cell structures. The cellular heterogeneity raised by tumor evolution presents another instance (5, 6). The copy number gain, neutral, and loss classify tumor cells into aneuploid, diploid, and hypodiploid groups, respectively. Fluctuations of copy numbers in focal genome regions further categorize tumor cells into amplification or deletion subtypes. The cell cycle is a rudimentary biological process for cell replications (7). Human cells undergo a cycle G1 - S - G2/M - G1 over a 24-hour period, thus the cycling cells have three flat phase labels (G1, S, and G2/M). In addition, the cycling cells have an order that records the pseudo time course in the G1, S, and G2/M phases. The orders and phase labels reflect a hierarchical structure.

The recent maturation of single-cell sequencing technologies offers opportunities to profile large-scale single cells for their transcriptomics (8), genomics (5), epigenomics (9), *etc*. These technologies have blossomed revolutionary insights into cellular functional diversity under the aegis of assigning cells with similar molecular characteristics to the same group (1, 2). However, most existing clustering tools generate flat cell group (10–14). Moreover, the periodic pseudo-time inference tools neglect the hierarchical structure of cycling cells (15–18). Neglection of the underlying nested structures of cells prevents full-scale detection of cellular functional diversity.

To address the issue, we incorporate *cell hierarchy* to illustrate the nested structure of cellular functional diversity. Cell hierarchy is a tree-like structure with multiple layers that capture cellular heterogeneity. From the root to the tips, the cellular heterogeneity decays. This study focuses on four main layers: global, subpopulation, club, and cell. The global layer is the root that exemplifies the whole cell population, e.g., immune cells. In contrast, the cell groups in the second and third main layers resemble *cell subpopulations* and *cell clubs*, respectively. The cell subpopulation is a broad category of cells, such as B cells and T cells (4). Cell clubs within one cell subpopulation catalog the cellular heterogeneity in a finer resolution; that is, the cells share high functional similarity within a single cell club. For example, T cell subpopulation owns helper and cytotoxic T cell clubs (4). The tip layer holds individual cells carrying *cell orders*, which signify the dynamic nuance of cell changes within a cell club, e.g., cellular heterogeneity varies along a periodic time course for cells undergoing a cycling process (7).

The actual cell hierarchy is difficult to determine; here, we develop SEAT, Structure Entropy hierArchy deTection, to build a pseudo cell hierarchy leveraging structure entropy to characterize the nested structures in cell-cell graphs. Structural entropy has been proposed in structural information theory to measure the dynamic global uncertainty of complex networks (19), and has benefited several biological fields (20–24). SEAT constructs cell hierarchies using a full-dimensional or dimensionally reduced single-cell molecular profile as input, and delivers the global-subpopulation-club-cell layers from the hierarchies. We apply SEAT to 36 datasets that cover single-cell RNA (scRNA), single-cell DNA (scDNA), single-cell assay for transposase-accessible chromatin (scATAC), and scRNA-scATAC multiome. SEAT detects the functional diversity of these single-cell omics data with cell hierarchy from three perspectives: cell subpopulation detection, cell club investigation, and periodic cell cycle pseudo-time inference.

Visualizing the functional diversity of single cells is essential since visual inspection is the most direct approach to studying the structure and pattern of cells. Nonlinear dimension reduction is a trending visualization method for high-dimensional biological data (25). Nevertheless, state-of-the-art single-cell visualization tools neglect the nested structure of cells by merely capturing at most two levels (global or local) of cell patterns (26–28). To tackle the issue, SEAT provides a component to embed the cells into a low-dimensional space by incorporating the multiple layers from the cell hierarchy as prior knowledge. Experiments demonstrate that SEAT consistently visualizes the hierarchical layout of these cells in the two-dimensional space for the above single-cell datasets.

## Method

### Problem formulation

#### Constructing cell-cell similarity graph

For a single-cell molecular data tabulated in a matrix, columns and rows refer to cells and their molecular features. For instance, the feature can be a gene or genome region. An entry in the matrix measures the value of the corresponding cell-feature pair, e.g., gene expression, copy number variation, or chromatin accessibility.

We reduce the dimensionality of the single-cell molecular matrix to a low-dimensional matrix ***X*** to mitigate the curse of dimensionality. We construct a dense cell-cell similarity graph *G* = (*V, E*) with Gaussian kernel 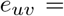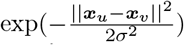 with *σ* as standard deviation of ***X***. Edge weight *e_uv_* stands for the similarity between cells *u* and *v* in graph *G*.

#### Hierarchical coding tree

A coding tree *T* of a cell-cell graph *G* = (*V, E*) is a hierarchical multi-nary partitioning of the cell set *V*, preserving the nested information in *G*. For clarity, we use *u* and *v* to represent the cells and *μ* and *ν* to represent tree nodes. Each tree node *μ* ∈ *T* codes a cell subset *U* ⊂ *V*. Denote the cell set coded by a node *μ* ∈ *T* as *V* (*μ*). The root node *r* codes *V* and node *μ* codes *U*, i.e., *V* (*r*) = *V* and *V* (*μ*) = *U*. Denote the children of *μ* as *C*(*μ*). The children nodes *C*(*μ*) of the tree node *μ* ∈ *T* partition the cells represented by *μ*; that is, 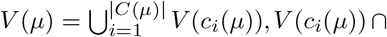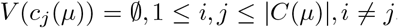, where *c_i_*(*μ*) signifies the *i*-th child node of *μ* and | · | denotes cardinality. A leaf node *t* codes one or multiple cells with a specific order *π*(*t*) ∈ ℕ^|*V* (*t*)|^. For each cell *u* ∈ *V* there is a unique leaf node *t* ∈ *T* such that {*u*} ⊆ *V* (*t*).

#### Coding tree represents the hierarchy of subpopulations, clubs, and cells

Given a pool of cells *V* which own *k* cell subpopulations, an ideal coding tree *T* holds *k* disjoint sub-trees rooted at nodes Λ = {*λ*_1_, …, *λ_k_*} which encode *k* cell sets 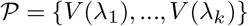 that match the cell subpopulations. Denote the subtree *T_λ_* ⋐ *T* rooted at *λ* as *subpopulation tree*. Suppose *T_λ_* has *ℓ_λ_* leaves 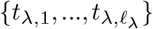, they encode *ℓ_λ_* cell sets 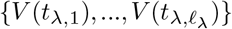 that represent cell clubs inside cell subpopulation *V* (*λ*) in a finer resolution; that is, the cells share high similarity inside one cell subpopulation. In coding tree *T*, the total *ℓ* leaves signify the *ℓ* cell clubs 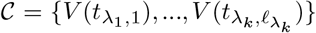. Moreover, as cells in each cell club *t* has a specific order *π*(*t*) ∈ ℕ^|*V* (*t*)|^, the ideal coding tree *T* also presents an overall cell order 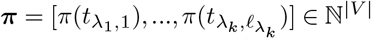 according to the order of leaves from left to right.

Determining the hierarchy of subpopulations, clubs, and cells is now a hierarchical coding tree construction problem - partitioning the graph *G* hierarchically to optimize a metric. In this work, the metric is the global dynamical complexity of the graph measured by structure entropy (19–24).

#### Measuring coding tree with structure entropy

Recall *e_uv_* is the edge weight between cells *u* and *v* for *G*. Term the volume of *μ* ∈ *T* as the sum of degrees of all cells in *V* (*μ*), *vol*(*μ*) = ∑_*u*∈*V* (*μ*),*v*∈*V*_ *e*_*uv*_. Define *g*(*μ*) as the total weights of edges from cells in *V* (*μ*) to *V* − *V* (*μ*), *g*(*μ*) = ∑_*u*∈*V* (*μ*),*v*∈*V* − *V* (*μ*)_ *e_uv_*. If *μ* ≠ *r*, its structure entropy is

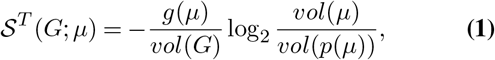

where *p*(*μ*) is the parent node of *μ*, *vol*(*G*) = ∑_*u*,*v*∈*V*_ *e_uv_* is the sum of all the edges in the graph, thus *vol*(*G*) = *vol*(*r*) signifies the volume of the whole graph or the root *r*. The root *r* has structure entropy 0; that is, 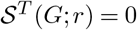.

Denote *t*(*u*) as the leaf node where cell *u* belongs to, the structure entropy of cell *u* in *T* is

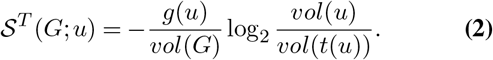

The structure entropy of graph *G* coded by tree *T* is the sum of the structure entropy of all tree nodes and all cells,

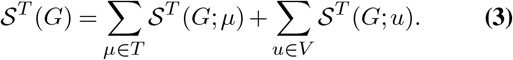

An ideal coding tree *T* captures the optimal hierarchy of subpopulations, clubs, and cells. Finding the optimal coding tree *T* for the graph *G* is to find the minimum structure entropy 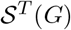 which diminishes the global variance at the random walk of *G* to a minimum.

#### Algorithm of SEAT

In previous work, we have proven that for a graph *G*, there exists a binary hierarchy of minimum structure entropy (23). Thus, SEAT searches the ideal coding tree *T* from the binary hierarchies (Fig. 1A). We first construct a sparse graph *G_s_* from dense graph *G*, then form cell club hierarchies with minimal structure entropy 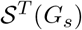 from sparse graph *G_s_* with agglomerative and divisive heuristics. Then, we search the cell subpopulations by optimizing the structure entropy of the dense graph *G* constrained by the heuristic hierarchies. Finally, we embed the graph *G* into a low-dimensional space by adding the global-subpopulation-club layer constraints from cell hierarchy *T*.

**Fig. 1.**
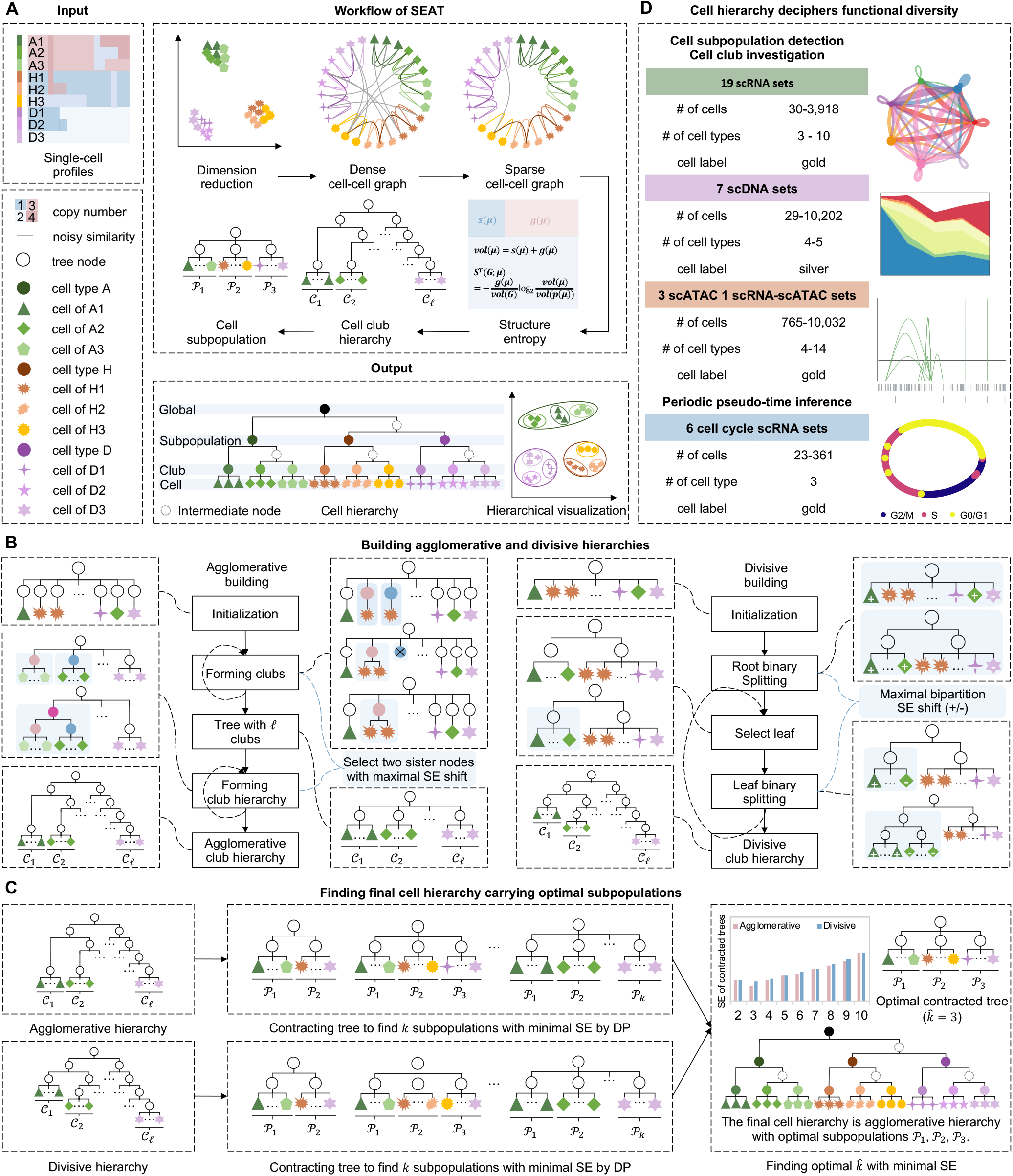
The schematic overview of SEAT. **A** The workflow of SEAT. **B** The algorithm of agglomerative and divisive hierarchy building. **C** The algorithm of finding the final cell hierarchy carrying optimal subpopulations. **D** The summary of experimental settings.

#### Graph sparsification

We sparsify the dense graph *G* with k-nearest neighbors (kNNs), resulting in a sparse graph *G_s_* = (*V, E_s_*) with a binary edge weight. If cell *u* is the k-nearest neighbor of cell *v* or cell *v* is the k-nearest neighbor of cell *u* in original graph *G*, *e_uv_* = 1; otherwise *e_uv_* = 0.

#### Building cell club hierarchy

With the sparse graph *G_s_*, we form cell club hierarchies with minimal structure entropy 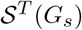 with agglomerative and divisive heuristics (Fig. 1B).

##### Agglomerative hierarchy building

The agglomerative hierarchy building consists of three steps: initialization, forming clubs, and building club hierarchy.

We initialize the tree of height one, the root node *r* has |*V*| immediate children, where each child node *t* is a leaf node that covers a single cell of *u*, *V* (*t*) = {*u*}. The initialized tree is multi-nary.

We merge the leaf nodes repeatedly to form cell clubs. A leaf has one of the two possible statuses at each iteration, individual or merged. Initially, all the leaves are labeled as individual. Two tree nodes *μ* and *ν* are referred to connected if there are inter-node edges between *V* (*μ*) and *V* (*ν*) in sparse graph *G_s_*. We merge an individual leaf *μ* with its connected sister *ν* by extracting *μ* and *ν* from *T* and creating a new node *μ*′ which codes all cells in *V* (*μ*) and *V* (*ν*). The new node *μ*′ is a child of root and a leaf labeled as merged. The pair (*μ, ν*) is chosen by the largest merging structure entropy change 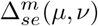 (Supplementary Methods). This merging operation repeats until i) there is no more individual leaf connected to other sister leaves; or ii) there is no pair (*μ, ν*) yields a non-negative structure entropy difference. Then, all leaves are labeled individual, triggering subsequent iterations of the merging procedure until no non-negative structure entropy shift is possible. The above will lead to a multi-nary coding tree *T* of a height of one and *ℓ* leaves. We assume each leaf presents a cell club, and the cell order is the merging order. To form the binary hierarchy of clubs, we iteratively combine sister node pair (*μ, ν*) of the root by inserting a new node *ω* as a child of the root and parent of *μ* and *ν*. The selection of (*μ, ν*) is guided by connectivity and the largest combining structure entropy change 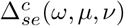 (Supplementary Methods). The combining operation repeats until the hierarchy is a binary coding tree.

##### Divisive hierarchy building

The second approach is to build the club hierarchy divisively. We initialize the tree with the root node *r* that codes all cells. The initialized tree has a zero height, with one node as both root and leaf. To form the hierarchy, we repeatedly split the leaf node *t* ∈ *T* into two children guided by maximizing the bipartition structure entropy change 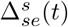. The solution of leaf split is the Fielder vector of the normalized graph Laplacian if the sparse graph *G_s_* is regular (Supplementary Methods). Thus, we heuristically obtain the bipartition according to the sign of values in Fielder vector (29), the cells with smaller Fielder vectors are placed on the left. The split stops if leaf node contains only two cells or 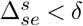, we set cutoff *δ* = 0.05. We assume that each leaf presents a cell club, and the value of Fielder vector reflects the cell order. Finally, we end up with a binary hierarchy *T* with *ℓ* clubs.

#### Finding cell subpopulations

Recall that an ideal coding tree *T* holds *k* disjoint subpopulation trees rooted at nodes Λ = {*λ*_1_, …, *λ_k_*} which encode *k* cell sets 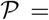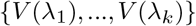 that match the cell subpopulations. To find the *k* subpopulations, we *contract* the heuristic club hierarchy *T* into a multi-nary tree 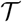 with a height of one (Fig. 1C). The contracted tree 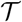 has a root node *r* holding *k* leaf children. Each leaf node 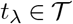 maps to a subpopulation tree *T_λ_* ⋐ *T* rooted at *λ*, thus *t_λ_* codes the cells from *T_λ_*, *p*(*t_λ_*) = *r*, *V* (*t_λ_*) = *V* (*λ*).

Given the heuristic club hierarchy *T*, contracting is optimized by minimizing the structure entropy 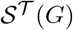 from dense graph *G*. The structure entropy associated with contracted tree 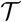 with *k* leaves focuses on measuring the global variance at the random walk of a dense graph *G* among *k* subpopulations, other than the variance in a finer cell-club resolution,

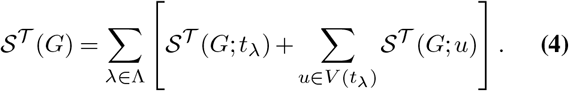

To minimize 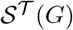, we adopt a recursive objective 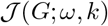 alongside the club agglomerative or divisive hierarchy *T*. Assume tree node *ω* in *T* has left and right children *μ* and *ν*, respectively. Finding *k* optimal subpopulation trees inside subtree *T_ω_* ⋐ *T* rooted at *ω* with minimum 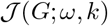 is equivalent to finding *k*′ and *k* − *k*′ subpopulation trees inside subtrees *T_μ_* ⋐ *T* and *T_ν_* ⋐ *T* rooted at *μ* and *ν* such that sum of structure entropy in the contracted tree 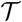 is minimal,

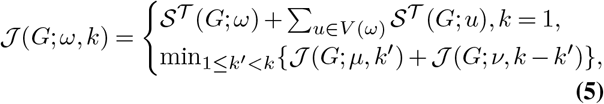

where *k* = 1 means *ω* is the root node of one subpopulation tree, which maps to one leaf node of the contracted tree 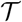.

We solve the contracting objective using dynamic programming. We record the minimal structure entropy 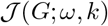 for finding *k* optimal subpopulations in a bottom-up way; that is, calculating from leaves to root. We trace back recursively to obtain the optimal cut-off *k*′ for each node starting from the root. If 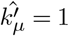 for one left child or 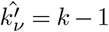 for a certain right child at that state, one subpopulation *V* (*μ*) or *V* (*ν*) is found (Supplementary Methods). In this way, we obtain the contracted tree 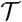 with *k* leaves representing *k* cell subpopulations.

#### Finding final cell hierarchy carrying optimal subpopulations

For 1 ≤ *k* ≤ *K* where *K* is constant number, the optimal 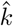 associated with the minimal structure entropy is the optimal cut-off *k*′ for root node, 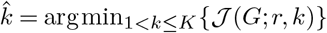.

The agglomerative and divisive hierarchies might have different hierarchical structures. The optimal subpopulations are subpopulations with less structure entropy (Fig. 1C and Supplementary Methods). We choose the cell hierarchy carrying optimal subpopulations as the final cell hierarchy.

#### Obtaining and visualizing cell order

We find the cell hierarchy *T* by minimizing the structure entropy of the sparse cell-cell graph. Given the cell hierarchy *T*, we obtain the cell order ***π*** ∈ ℝ^|*V* |^ with an in-order traversal and visualize the cell order periodically with an oval shape (Supplementary Methods).

#### Hierarchical visualization

To convert the cell-cell similarity graph *G* into *d*-dimensional latent space ***Y*** ∈ ℝ^*n*×*d*^ for visualization, state-of-the-art tool UMAP (26) adopts a cross-entropy (CE) objective,

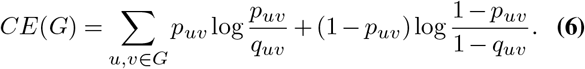

Here, *p_uv_* and *q_uv_* signifies the similarity of cells *u* and *v* in original graph *G* and the latent space, respectively. *q_uv_* is smoothly approximated by 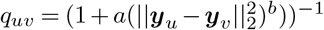, where *a* and *b* are constrained by a hyper-parameter *min-dist*, the effective minimum distance between cells in latent space. In this study, we adjust the above embedding strategy by incorporating the final cell hierarchy. Recall that the cell partition and 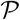 and 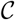 correspond to the *k* and *ℓ* cell subpopulations and clubs, respectively. Assume cell partition 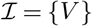 contains the one global cell population. Based on the cell partition 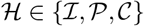, we assign the inter-connections between different cell groups to zero, resulting in a graph 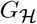 that focuses on the cell-cell similarity inside one cell group. We minimize the disparity of cell-cell similarity between the embedding space and 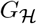 with the objective

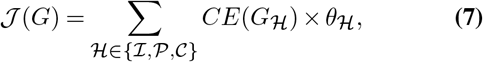

where hyper-parameters 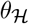 are the training weights of different cell partition resolutions obtained from cell hierarchy. We initialize the low-dimensional embedding ***Y*** with graph Laplacian (30) of 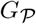, make *min-dist* equals 0.1, set 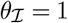, 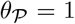, 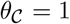, and minimize 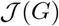 to convergence with Adam gradient descent.

#### Outlier detection

Cellular abnormalities may distort the entire cell hierarchy, thus affecting the efficacy of cell subpopulation and club detection, cell cycle pseudo-time inference, and hierarchical visualization. Thus, we have implemented the average kNN outlier detection. We calculate the mean distance ***d*** ∈ ℝ^*n*^ given the single-cell molecular representation of *n* cells. *d_i_* is the mean distance of *i*-th cell to its k-nearest neighbors. By default, we consider the cell with an average kNN distance *d* exceeding a distance cutoff 0.5 as the outlier. We also provide a distance percentile cutoff strategy, we regard the cell with an average kNN distance *d* surpassing a distance percentile cutoff (e.g. 95th percentile) as an outlier. The detected outliers will be assigned to label-1 and excluded from the cell hierarchy building.

#### Time complexity of SEAT

Under the graph *G* with *n* cells, the time complexity of SEAT is *O*(*n* log *n*) (Supplementary Methods).

### Experiment Setting

#### scRNA data

We collect nineteen scRNA datasets with gold standard cell type labels (31–43), the description of the datasets and the download links are in Supplementary Table S1 and Supplementary Method. For these scRNA datasets, the dimension reduction transformer is UMAP (26). We adopt Seurat “FindAllMarkers” function (44) for differential expression analysis. The log2 fold change, log2(FC), of the average expression between two groups is measured. The fold change significance p-value is evaluated by the Wilcoxon Rank Sum test, and the adjusted p-value is calculated with Bonferroni correction. The filtering criteria are log2(FC) ≥ 0.25, p-value < 0.05, and adjusted p-value < 0.05. Cell-cell communication analysis is conducted with CellChat (45) with default database and parameters. Any ligand-receptor interaction with less than ten supporting cells is filtered.

We also collect six scRNA datasets with gold standard cell cycle labels (Supplementary Table S2). Dataset H1-hESC has 247 human embryonic stem cells (hESCs) in G0/G1, S, or G2/M phases identified by fluorescent ubiquitination-based cell cycle indicators (46). The count expression profile and cell cycle labels are obtained with accession code GSE64016. Datasets mESC-Quartz and mESC-SMARTer have 23 and 288 mouse embryonic stem cells (mESCs) sequenced by Quartz-seq and SMARTer, respectively (47, 48). Their G0/G1, S, and G2/M phases are labeled by Hoechst staining. The count expression profiles and cell cycle labels are obtained with accession codes GSE42268 and E-MTAB-2805. Datasets 3Line-qPCR_H9, 3Line-qPCR_MB, and 3Line-qPCR_PC3 own 227 H9 cells, 342 MB cells, and 361 PC3 cells, respectively. The cell cycle stages G0/G1, S, and G2/M are marked by Hoechst staining (32). The raw log2 count expression profiles and cell labels are from the paper’s dataset S2. The imputation and dimension reduction are conducted by SMURF (49) and UMAP (26). We adopt Seurat (44) for differential expression analysis as described above. Cell-cell communication analysis is conducted with CellChat (45) with default database and parameters. Any ligand-receptor interaction with less than ten supporting cells is filtered. Gene Ontology (GO) is performed with ShinyGO 0.76 (50).

#### scDNA data

We collect seven scDNA datasets (Supplementary Table S1). Navin_T10 contains 100 cells from a genetically heterogeneous (polygenetic) triple-negative breast cancer primary lesion T10, including five cell subpopulations: diploid (D), hypodiploid (H), aneuploid 1 (A1), aneuploid 2 (A2), and pseudo-diploid (P) (51). Navin_T16 holds 52 cells from genetically homogeneous (monogenetic) breast cancer primary lesion T16P and 48 cells from its liver metastasis T16M, including four cell subpopulations: diploid (D), primary aneuploid (PA), metastasis aneuploid (MA), and pseudo-diploid (P) (51). The Ginkgo copy number variation (CNV) profiles of Navin_T10 and Navin_T16 are downloaded from http://qb.cshl.edu/ginkgo (52). The silver standard array comparative genomic hybridization (aCGH) data of Navin_T10 and Navin_T16 are downloaded with GEO accession code GSE16607 (53).

Dataset 10x_breast_S0 is a large-scale 10x scDNA-seq set without known cell population labels, where 10,202 cells from five adjacent tumor dissections (A, B, C, D, and E) of triple-negative breast cancer are sequenced. The Bam files are downloaded from 10x official site https://www.10xgenomics.com/resources/datasets. We inferred the total CNV profile utilizing Chisel (54).

Ni_CTC sequenced 29 circulating tumor cells (CTCs) across seven lung cancer patients (55). McConnel_neuron profiles 110 cells from human frontal cortex neurons, with an extensive level of mosaic CNV gains and losses (56). Lu_sperm sequenced 99 sperm cells with chrX-bearing, chrY-bearing, and aneuploid groups (57). Wang_sperm performed single-cell sequencing on 31 sperm cells with CNV gains and losses (58). The Ginkgo CNV profiles of these datasets are downloaded from http://qb.cshl.edu/ginkgo (52).

#### scATAC and scRNA-scATAC multiome data

We collect three public scATAC-seq data as benchmarking sets with gold standard cell type labels (Supplementary Table S1). scatac_6cl is a mixture of six cell lines (BJ, GM12878, H1-ESC, HL60, K562, and TF1) with 1224 cells (59). Hematopoiesis owns 2210 single-cell chromatin accessibility profiles from eight human hematopoiesis cell subpopulations (CLP, CMP, GMP, HSC, LMPP, MEP, MPP, and pDC) (60). T-cell composes of four T-cell subtypes (Jurkat_T_cell, Naive_T_cell, Memory_T_cell, and Th17_T_cell) with a total of 765 cells (61). We collect a multiome of scRNA and scATAC dataset PBMC (human peripheral blood mononuclear cells) with 10,032 cells across fourteen cell types.

We downloaded the scOpen (62) processed accessibility profiles and cell labels from https://github.com/CostaLab/scopen-reproducibility. UMAP (26) embedded data are used to construct the kNN graphs for each dataset. We adopt Cicero (63) to explore the dynamically accessible element status in different scatac_6cl GM12878 cell clubs.

#### Evaluating cell subpopulation detection

To detect cell subpopulations, some clustering methods require the number of clusters prespecified, while others can determine the number of clusters automatically. The SEAT package supports both. Our package requires no prespecified number of clusters by default, that is, SEAT(sub). If the number of clusters required is *k*, we denote the method as SEAT(k). When the context is clear, we refer to them as predefined-k and auto-k modes, respectively.

In the predefined-k mode, we access the clustering accuracy of SEAT agglomerative hierarchy and divisive hierarchy with predefined cluster number *k* given by the actual number of ground truth cell types, namely Agglo(k) and Divisive(k). We regard the clustering result with a lower structure entropy from agglomerative and divisive hierarchies as SEAT(k). Baselines are hierarchical clustering (HC) with four linkage strategies (ward, complete, average, and single) (12), K-means (11), and spectral clustering (10). We run them with default parameters. As the leading tool for single-cell clustering, Louvain (13) and Leiden (14) automatically detect how many communities are inside the cell-cell similarity graph. They obtain different numbers of communities at various resolutions. To benchmark Leiden and Louvain in the predefined-k setting, namely Leiden(k) and Louvain(k), we heuristically adjusted the resolution 20 times to see if the number of communities was the same as the predefined cluster number *k*.

As the predefined *k* is undetermined in most real-world scenarios, we evaluate the auto-k clustering efficacy of SEAT cell hierarchy, agglomeration hierarchy, and divisive hierarchy, namely SEAT(sub), Agglo(sub), and Divisive(club). The baselines are Leiden and Louvain with default parameters. We also assess the clustering obtained from agglomerative and divisive hierarchy clubs, namely Agglo(club) and Divisive(club).

Adjusted Rand index (ARI) (64) and adjusted mutual information (AMI) (65) are adopted as clustering accuracy. They measure the concordance between clustering results and ground truth cell types. Perfect clustering has a value of 1, while random clustering has a value less than or near 0.

#### Evaluating cell cycle pseudo-time inference

SEAT cell hierarchy, agglomerative hierarchy, and divisive hierarchy generate cell orders representing the cell cycle pseudo-time for scRNA data, namely, SEAT(order), Agglo(order), and Divisive(order). We access the pseudo-time inference accuracy of SEAT given by the actual order of ground truth cell cycle phases. Benchmark methods are hierarchical clustering (HC) with four linkage strategies (ward, complete, average, and single) (12), since an in-order traversal of HC hierarchies also generates cell orders. Furthermore, we benchmark our method with four state-of-the-art tools predicting the cell cycle pseudo-time, CYCLOPS (15), Cyclum (16), reCAT (17), and CCPE (18). We run them with default parameters. CCPE fails the tasks when we follow its GitHub instruction, so we exclude CCPE for final comparison.

The change index (CI) is used to quantitatively assess the accuracy of cell pseudo-time order against known cell cycle phase labels (17). An ideal cell order changes label *k* − 1 times, where *k* = 3 is the ground truth cell cycle phase number. The change index is defined as 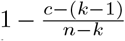, where *c* counts the frequency of label alters between two adjacent cells, and *n* is the number of cells. A value of 0 suggests the cell order is utterly wrong with *c* = *n* − 1, while 1 indicates a complete match between cell order and ground truth cell cycle phase with *c* = *k* − 1.

#### Evaluating hierarchical visualization

We evaluate the effiacy of SEAT hierarchical visualization, SEAT(viz), with state-of-the-art visualization tools UMAP (26), TSNE (27), and PHATE (28). The dense cell-cell similarity graph *G* is used as input, UMAP, TSNE, and PHATE are run with default parameters.

#### Evaluating cell outlier detection

We simulate the gene expression profiles of 500 cells with five subpopulations using Splatter (66). We randomly produce 20 cell outliers with gene expression disparting from all five subpopulations. We evaluate SEAT cell subpopulation detection i) with and without the average kNN outlier detection; ii) with different combinations of parameters (nearest neighbor number, distance cutoff, and distance percentile cutoff). The outliers are considered as a distinct group, thus the ARI and AMI are used to measure the clustering accuracy.

## Results

### Overview of SEAT

SEAT builds a cell hierarchy annotated with global-subpopulation-club-cell layers computationally from single-cell data (Fig. 1). First, SEAT constructs a pair of dense and sparse cell-cell similarity graphs with a full-dimensiona or dimensionally reduced single-cell molecular profile as input (Fig. 1 A). Second, we detect cell clubs, determine the order of cells within each cell club, and build the pseudo club hierarchies by minimizing the structure entropy of the sparse graph with agglomerative (Agglo) and divisive (Divisive) heuristics (Fig. 1B, Methods). We term the cell clubs and orders derived from agglomerative and divisive hierarchies as Agglo(club), Agglo(order), Divisive(club), and Divisive(order). Next, we use dynamic programming to find optimal subpopulations from agglomerative and divisive hierarchies, namely, Agglo(sub) and Divisive(sub). We choose the hierarchy carrying the lower subpopulation structure entropy as the final cell hierarchy (Fig. 1C, Methods). Hence, SEAT outputs the final cell hierarchy carrying with subpopulations, clubs, and orders, namely, SEAT(sub), SEAT(club), and SEAT(order) (Fig. 1A). Furthermore, by incorporating hierarchical cell partition layers, SEAT provides a component, SEAT(viz), to embed cells into a low-dimensional space while preserving their nested structures for improved visualization and interpretation (Fig. 1A).

### Cell hierarchy catalogs functional diversity at the subpopulation and club level from scRNA data

We have applied SEAT to nineteen scRNA datasets carrying gold standard cell type labels. The first nine sets are cell line mixtures, including p3cl (31), 3Line-qPCR (32), sc_10x, sc_celseq2, sc_dropseq, sc_10x_5cl, sc_celseq2_5cl_p1, sc_celseq2_5cl_p2, and sc_celseq2_5cl_p3 (33). We have four datasets Yan (34), Deng (35), Baise (36), and Goolam (37) which sequence single cells from human or mouse embryos at different stages of development (zygote, 2-cell, early 2-cell, mid 2-cell, late 2-cell, 4-cell, 8-cell, 16-cell, 32-cell, early blast, mid blast, and late blast). The last six datasets are Koh (38), Kumar (39), Trapnell (40), Blakeley (41), Kolodziejczyk (42), and Xin (43), which profile different cell types in single-cell resolution. To access the efficacy of SEAT in cell subpopulations detection, we utilize the adjusted rand index (ARI) (64) and adjusted mutual information (AMI) (65) as clustering accuracy and benchmark SEAT with state-of-the-art clustering tools (spectral clustering (10), K-means (11), hierarchical clustering (12), Louvain (13), and Leiden (14)) with predefined-k and auto-k modes (Methods, Supplementary Fig. S1-S3). In predefined-k mode, SEAT(k) demonstrates comparable or higher clustering accuracy compared to other clustering baselines on most datasets (Fig. 2A). Notably, Louvain(k) and Leiden(k) are unable to generate a clustering that exactly matches the number of ground truth labels after 20 different resolution trials for the Goolam and Kolodziejczyk (Fig. 2A and Supplementary Fig. S2). Under the auto-k mode, SEAT(sub) outperforms Louvain and Leiden on all nineteen sets. The clustering accuracies of SEAT(sub) are comparable to or better than the best clustering results with predefined-k clustering tools with the ground truth cluster number provided. This is attributed to the fact that SEAT(sub) finds a cluster number close to the ground truth (Fig. 2 B). Louvain and Leiden have the lowest clustering accuracy because they prefer more clusters. The two-dimensional data embedded by UMAP from full-dimensional single-cell expression profiles are inputs of all clustering tools; and the visualizations of them show that the ground truth labels are mixed for the majority of datasets (Supplementary Fig. S4-S5), explaining the low clustering accuracy of both predefined-k and auto-k clustering tools.

**Fig. 2.**
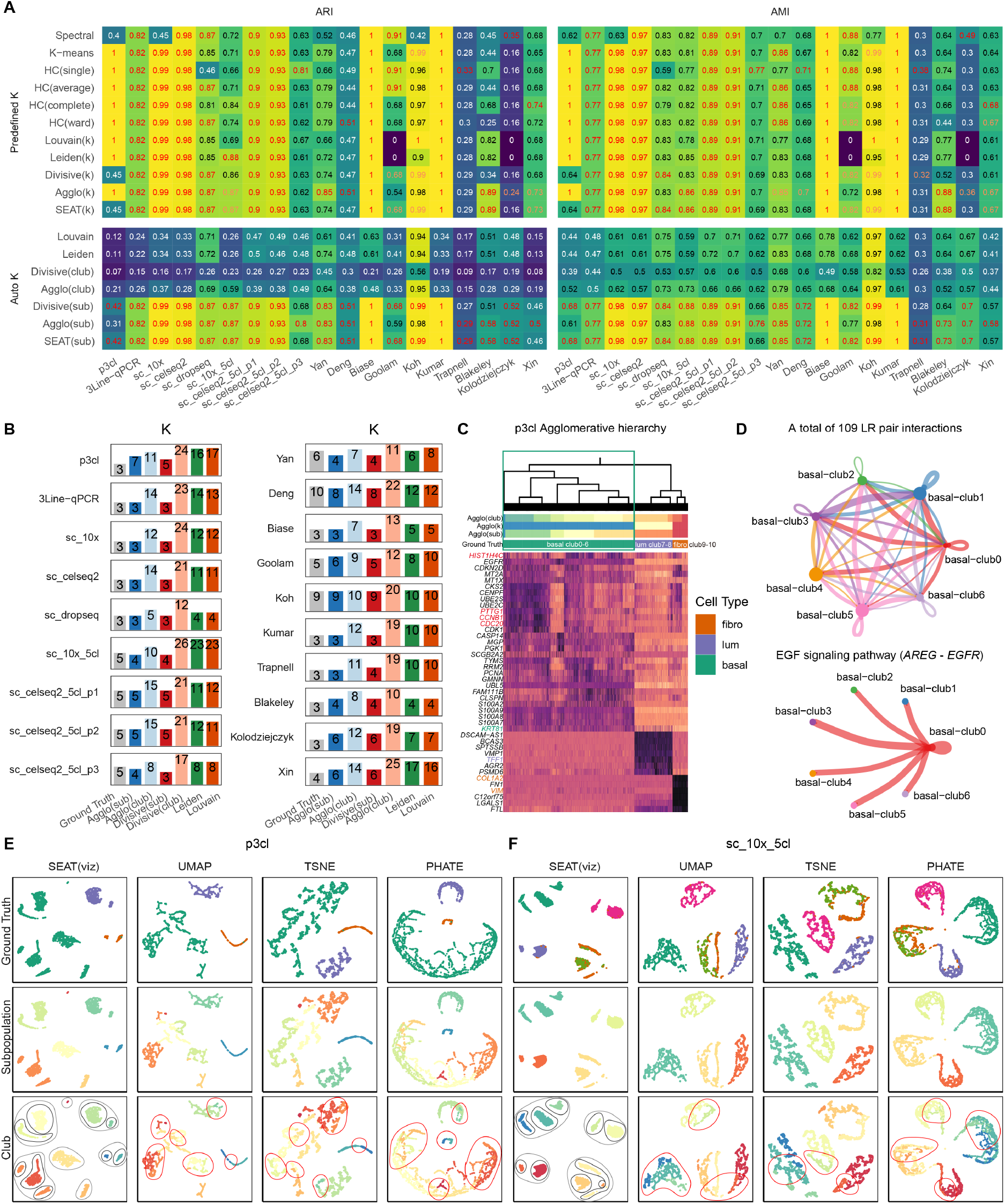
Applying SEAT on nineteen scRNA datasets. **A**. The adjusted rand index (ARI) and adjusted mutual information (AMI) of predefined-k and auto-k clustering tools. The best scores are colored red for each dataset in predefined and auto clustering benchmarking separately. If SEAT gets second place, we color the score orange. Spectral: spectral clustering. HC(single), HC(average), HC(complete), and HC(ward): hierarchical clustering with single, average, complete, and ward linkage. Louvain(k) and Leiden(k): Louvain and Leiden in predefined-k mode. Divisive(k) and Agglo(k): the cell subpopulations from divisive and agglomerative hierarchy in predefined-k mode. SEAT(k): the cell subpopulations from SEAT cell hierarchy in predefined-k mode. Divisive(club) and Agglo(club): the cell clubs from the divisive and agglomerative hierarchy. Divisive(sub) and Agglo(sub): the cell subpopulations from divisive and agglomerative hierarchy in auto-k mode. SEAT(sub): the optimal subpopulations from SEAT cell hierarchy in auto-k mode. **B**. The number of subpopulations detected for auto-k clustering tools. **C**. The top five differentially expressed genes in agglomerative hierarchy clubs for p3cl. **D**. The cell-cell communications among seven agglomerative hierarchy clubs for breast cancer basal-like epithelial cell line in p3cl. LR: ligand-receptor. **E**-**F** SEAT(viz), UMAP, TSNE, and PHATE plots for p3cl and sc_10x_5cl. The cells are colored with subpopulations, clubs, and ground truth. The gray and black circles in the SEAT(viz) plot indicate the subpopulation and club boundaries, respectively. In UMAP, TSNE, and PHATE plots, the red circles mark the unclearly segregated cell clubs. SEAT(viz): the hierarchical visualization from SEAT cell hierarchy.

SEAT offers hierarchical structures of cells to study cellular functional diversity. We leverage differential gene expressions to investigate the biological interpretations of these hierarchies. In Supplementary Fig. S6-S7, differentially expressed genes (*p* < 0.05) between cell hierarchy clubs reveal distinct patterns that match ground truth cell subpopulations. Furthermore, visible marker gene patterns reveal the functional diversity among cell clubs within one cell subpopulation. We focus on the top five differentially expressed genes for each dataset (Supplementary Fig. S8-S11). As the subpopulation detection accuracy of agglomerative hierarchy is 1 for p3cl dataset, we investigate the functional diversity revealed from the agglomerative hierarchy other than the divisive hierarchy. The agglomerative hierarchy revealed three cell subpopulations for p3cl, which correspond to the three ground truth cell types, basal (*KRT81*), luminal (*TFF1*), and fibroblast (*COL1A2* and *VIM*) (Fig. 2D). We observe that each of the basal, luminal, and fibroblast has two major subclasses, controlled by the expression of cell cycle genes (*HIST1H4C*, *CDC20*, *CCNB1*, and *PTTG1*). Cell-cell communication analysis finds a total of 109 significant (*p* < 0.05) ligand-receptor (LR) pair interactions among seven agglomerative hierarchy clubs for breast cancer basal-like epithelial cell line in p3cl. The LR interactions belong to nine signaling pathways AGRN, CD99, CDH, EGF, JAM, LAMININ, MK, NECTIN, and NOTCH (Fig. 2D and Supplementary Fig. S12). In particular, there is a distinct breast cancer cell club (basal-club0) that drives *AREG* -*EGFR*, an oncogenic signaling (67) in breast cancer, to all basal cells, resulting in a high level of *AREG* activated *EGFR* expression (Fig. 2E). The two cell clubs from the luminal subpopulation have six significant (*p* < 0.05) LR interactions involving MK, SEMA3, and CDH signaling pathways (Supplementary Fig. S13). The fibroblast has three significant (*p* < 0.05) LR interactions, including two signaling pathways FN1 and ncWNT (Supplementary Fig. S13). The cell club fibro-club10 release *WNT5B* and then bind *FZD7* from fibro-club9, consistent with the observation that ncWNT is the predominant signaling pathway in skin fibroblasts (45).

Visualizations of two-dimensional data by UMAP from full-dimensional single-cell expression profiles reveal a dense layout (Supplementary Fig. S4-S5). The ground truth cell subpopulations are indistinctly separated in some high clustering accuracy datasets, and the cell clubs are densely arranged in each subpopulation clump. Here, we check whether SEAT hierarchical visualization eliminates the dense layout of clubs. We use the cell-cell graph constructed by SEAT as input and execute SEAT(viz), UMAP, TSNE, and PHATE, independently. In Fig. 2E-F and Supplementary Fig. S14-S18, SEAT(viz), UMAP, TSNE, and PHATE separate the ground truth cell type for most datasets. It should be noted that the patterns from SEAT(viz), UMAP, TSNE, and PHATE also correspond to the subpopulation layer annotations, validating SEAT subpopulation finding efficacy. At the cell club level, SEAT(viz) show a clear layout of cell clumps that correspond to the cell hierarchy; each cell club owns a distinct clump, and the distance between clubs belonging to the same subpopulation is within proximity. Although UMAP, TSNE, and PHATE capture the local structures of the clubs, the cell clubs marked with red circles are unclearly segregated.

### Cell hierarchy deciphers periodic cell cycle pseudo-time from single-cell data

We collect six scRNA cell cycle datasets, H1-hESC (46), mESC-Quartz (47), mESC-SMARTer (48), 3Line-qPCR_H9, 3Line-qPCR_MB, and 3Line-qPCR_PC3 (32) with gold standard G0/G1, S, or G2/M stages and build the cell hierarchies (Supplementary Fig. S19). In predefined-k and auto-k clustering benchmarking (Supplementary Fig. S20), SEAT illustrates higher or comparable clustering accuracy in the six datasets. SEAT predicts the optimal number of clusters closest to ground truth three, while Leiden and Louvain generally predict more clusters than SEAT. Further investigation shows that ground truth labels are mixed or not distinctly separated in two-dimensional data derived by UMAP for all datasets (Supplementary Fig. S21), explaining the poor performance of 3Line-qPCR data. Likewise, hierarchical visualization plots depict nested layouts corresponding to the cell hierarchies in visualization refinement experiments (Supplementary Fig. S22).

If we order the cells in cycling progress, cells from the same phase should be lined up adjacently as they share higher similarity. Thus, the cell order obtained from an ideal hierarchy could present a periodic pseudo-time order for cell cycle data. We visualize the cell order periodically with an oval plot, the placements of the cells in the oval represent their pseudo-time in the cell cycle (Fig. 3A and Supplementary Fig. S23). We access the cell ordering accuracy with the change index (CI) (17), which computes how frequently the gold standard cell cycle phase labels switch along the cell order. The benchmark methods are four conventional HC strategies (12) that offer a cell order. We also recruit state-of-the-art tools dedicating to predict the cell cycle pseudo-time, CYCLOPS (15), Cyclum (16), reCAT (17), and CCPE (18). SEAT demonstrates the highest ordering accuracy for all datasets, except for 3Line-qPCR_PC3, where SEAT wins the top two (Fig. 3B). We exclude CCPE as it fails the tasks. In all, this suggests that cell hierarchy obtained from SEAT facilitates the cell cycle pseudo-time order inference.

**Fig. 3.**
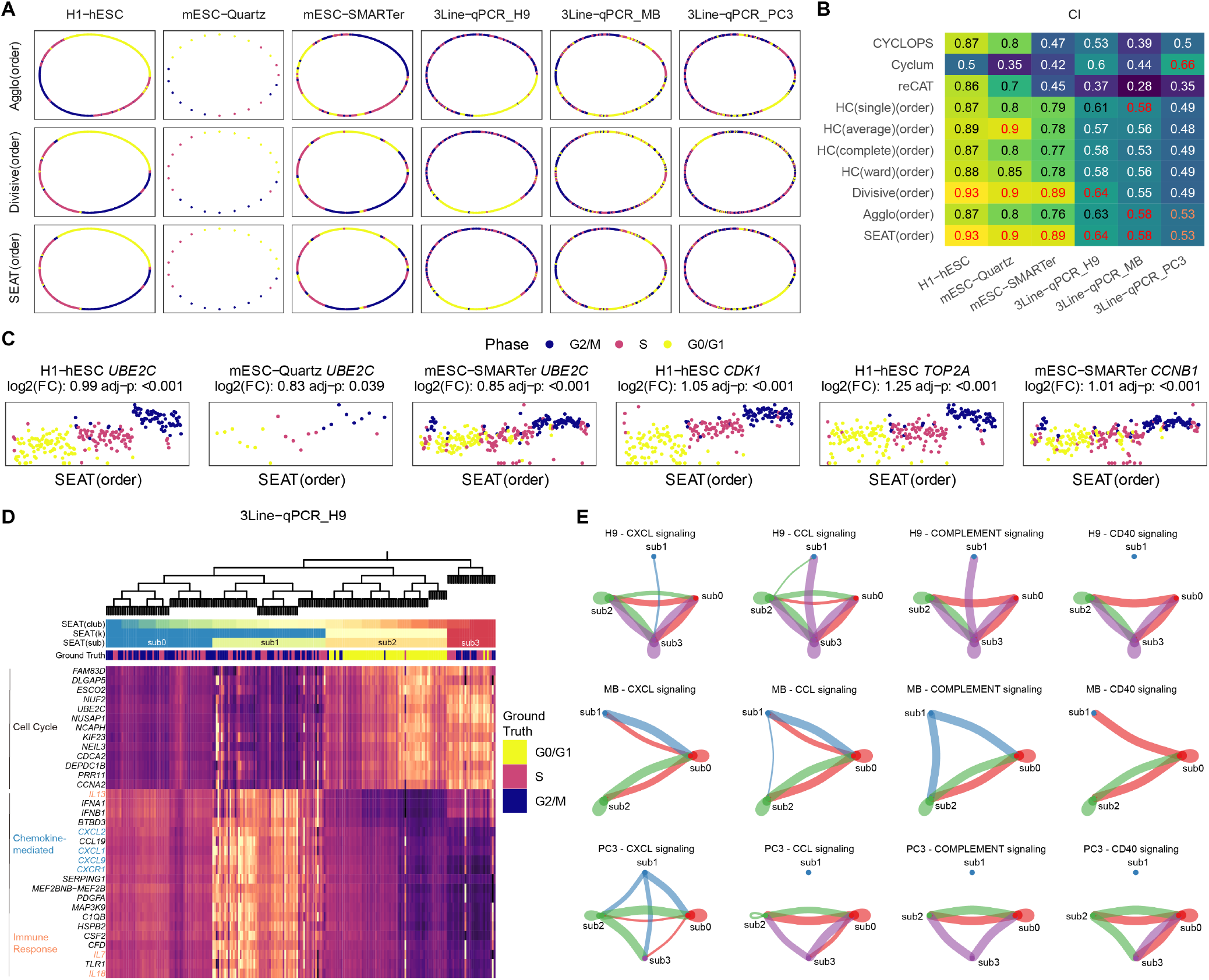
Applying SEAT on six scRNA cell cycle datasets. **A**. The oval visualization of cell pseudo-time. From left to right are H1-hESC, mESC-Quartz, mESC-SMARTer, 3Line-qPCR_H9, 3Line-qPCR_MB, and 3Line-qPCR_PC3. From top to bottom are cell orders obtained from agglomerative hierarchy, divisive hierarchy, and SEAT cell hierarchy; namely, Agglo(order), Divisive(order), and SEAT(order). **B**. The accuracy of cell pseudo-time order is measured by change index (CI) for baseline tools. The best scores are colored red for each dataset. If SEAT gets second place, we color the score orange. HC(single)(order), HC(average)(order), HC(complete)(order), and HC(ward)(order): the cell orders from hierarchical clustering with single, average, complete, and ward linkage. **C**. The normalized expression of M phase marker genes alongside the SEAT cell order. **D**. The top 20 differentially expressed genes in G0/G1, S, and G2/M ground truth phases for p3cl, arranged with SEAT cell hierarchy. SEAT(club): the cell clubs from SEAT cell hierarchy. SEAT(k): the cell subpopulations from SEAT cell hierarchy in predefined-k mode. SEAT(sub): the optimal subpopulations from SEAT cell hierarchy in auto-k mode. **E**. The cell-cell communications among SEAT cell subpopulations for H9, MB, and PC3 cell lines.

SEAT orders cells in H1-hESC, mESC-Quartz, and mESC-SMARTer alongside the oval that closely matches the G0/G1-S-G2/M cycle (Fig. 3A). Differential expression analysis among ground truth phases reveals distinct cell cycle phase markers (Supplementary Fig. S24). These visible cell cycle marker patterns remain consistent when rearranging with SEAT cell order (Supplementary Fig. S25). The top 20 differential expression genes (*p* < 0.05) for hESC and mESC cells include well-known cell cycle markers *UBE2C*, *TOP2A*, *CDK1*, and *CCNB1* (Supplementary Fig. S26). Their expressions rise progressively with SEAT recovered pseudo-time order and are peaked with significant fold changes at the M phase (Fig. 3C).

In H9, MB, and PC3 cell lines, the cell orders in the S and G2/M phases are partially arranged compared to the exact time course (Fig. 3A). The differential expression makers of ground truth phases show that there are sub-patterns within the S and G2/M phases. Moreover, there are similar patterns shared between the S and G2/M phases (Supplementary Fig. S24), suggesting the cause of poor performance in pseudo-time ordering. Interestingly, after rearranging the marker expression heatmap with SEAT cell hierarchy, we observe distinct marker gene patterns among SEAT discovered cell subpopulations (Supplementary Fig. S25). For the H9 cell line, SEAT detected four cell subpopulations (Fig. 3D), G0/G1 phase corresponds to sub2. Cell cycle S and G2/M phases together have three cell subpopulations, sub0, sub1, and sub3. The top 20 differential expression genes (*p* < 0.05) exhibits two groups (Fig. 3D). The genes from the first group are enriched in GO cell cycle signaling pathways. The genes from the second group are enriched in GO chemokine-mediated signaling and immune response pathways with CXC and IL gene families, respectively (Supplementary Fig. S27). We demonstrate the top 20 differential expression genes for MB and PC3 cell lines in Supplementary Fig. S26-S27. Finally, we verify the cellular interactions among cell subpopulations with cell-cell communication analysis. We find a total of 124, 87, and 77 significant (*p* < 0.05) LR pair interactions among cell subpopulations for H9, MB, and PC3 cell lines, respectively. All datasets exhibit CXCL, CCL, COMPLEMENT, and CD40 signaling interactions among cell subpopulations (Fig. 3E).

### Cell hierarchy detects rare subclones on scDNA data

SEAT catalogs the clonal subpopulations of solid tumors and circulating tumor cells in four scDNA datasets. SEAT also identifies the CNV substructures of neuron and gamete cells in three scDNA datasets. Owning to the unique characteristics of CNV profiles, we only adopt SEAT agglomerative hierarchy to investigate the functional diversity of CNV sub-structures.

Navin *et al*. have profiled 100 cells from a genetically heterogeneous (polygenetic) triple-negative breast cancer primary lesion Navin_T10 (51). Fluorescence-activated cell sorting (FACS) analysis has confirmed that Navin_T10 carried four main cell subpopulations: diploid (D), hypodiploid (H), aneuploid A (A1), and aneuploid B (A2). Furthermore, Navin *et al*. have reported pseudo-diploid cells (P) with varying degrees of chromosome gains and losses from diploids. They are unrelated to the three tumor cell subgroups (H, A1, and A2) (51). Therefore, given whole-genome single-cell CNV profiles as input, we verify whether SEAT and the state-of-the-art clustering tools identify the four major cell groups and the rare pseudo-diploid cell group (Fig.4A). In predefined-k mode, SEAT agglomerative hierarchy successfully recognizes five cell subpopulations consistent with the patterns of CNV profiles. From top to bottom, the ranks are cancer normal cell group (D), pseudo-diploid cell subgroups (P), sub-groups H, and two tumor aneuploid groups, A1 and A2 (Fig. 4A). Leiden(k) and Louvain(k) fail at this task after 20 different resolution trials. Four HC strategies and K-means fail to distinguish the four pseudo-diploid cells as in the Navin *et al*.’s HC trial (51). Spectral clustering performs poorly by mixing tumor and normal cells. Regarding auto-k clustering algorithms, agglomerative hierarchy identifies five concordant subpopulations as predefined-k mode. Leiden and Louvain fail with the same sparse cell-cell similarity graph as input. Then, we leverage CNV density signals detected by aCGH from FACS identified D, H, A1, and A2 dissections of T10 (53) as silver standard to validate the clustering result. We calculate the pairwise Spearman correlation and Euclidean distance (L2-norm) between scaled single-cell CNV profiles and aCGH CNV signals. As a proof of concept, the single-cell CNV profiles of three bottom clusters separately own higher correlation and lower distance to aCGH profiles of H, A1, and A2 sections. The cells in the up-permost subpopulation detected by SEAT have almost zero correlation and the lowest distance with aCGH D sections, suggesting that they are diploid cells. Pseudo-diploid cells illustrate a low correlation with all aCGH sections, validating their unique CNV profiles. Navin *et al*. have sequenced 100 cells from a monogenic triple-negative breast cancer tumor and its seeded liver metastasis (Navin_T16) (51). SEAT clusters the 100 samples into four distinct subpopulations (Supplementary Fig. S28). Two are primary and metastasis aneuploid cells, corresponding to the published population structure. Notably, SEAT catalogs diploid cells and pseudo-diploid cells while baseline tools failed.

**Fig. 4.**
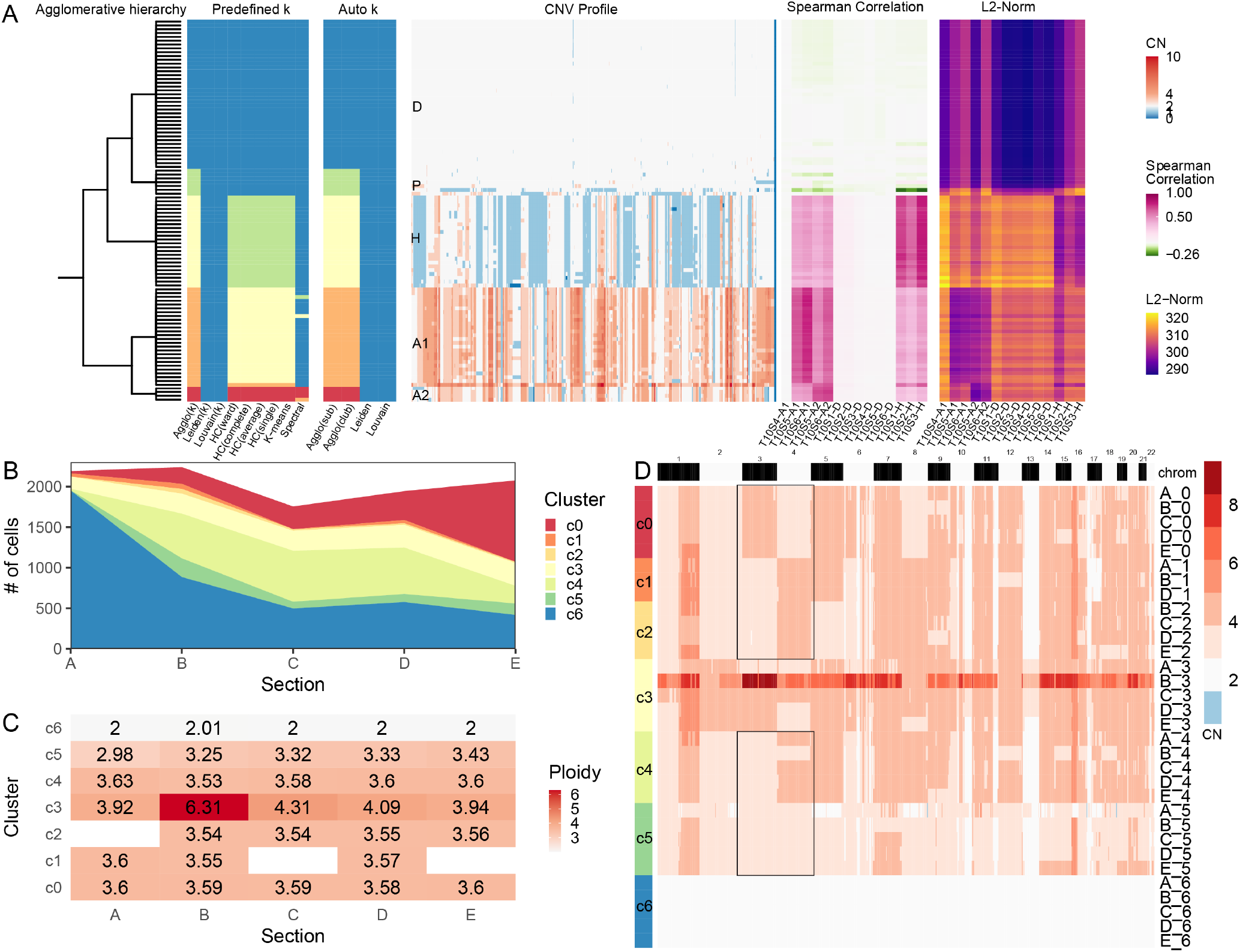
Applying SEAT on scDNA datasets. **A**. The analysis result of Navin_T10. From left to right is the SEAT agglomerative hierarchy, subpopulation detecting results for predefined-k (*k* = 5) and auto-k clustering tools, the whole genome single-cell CNV heatmap of T10, the Spearman correlation, and Euclidean distance (L2-Norm) between scaled copy number profiled by scDNA and copy number density profiled by aCGH. Spectral: spectral clustering. HC(single), HC(average), HC(complete), and HC(ward): hierarchical clustering with single, average, complete, and ward linkage. Louvain(k) and Leiden(k): Louvain and Leiden in predefined-k mode. Agglo(k): the cell subpopulations from agglomerative hierarchy in predefined-k mode. Agglo(club): the cell clubs from the agglomerative hierarchy. Agglo(sub): the cell subpopulations from agglomerative hierarchy in auto-k mode. **B**. The stacked area plot illustrates the SEAT subpopulations across 10x_breast_S0 tumor sections. Cluster c6 (blue) signifies the diploid cells. **C**. The mean ploidy of SEAT subpopulations across 10x_breast_S0 tumor sections. **D**. The whole-genome single-cell CNV heatmap of SEAT subpopulations across 10x_breast_S0 tumor sections. The black boxes highlight the mutually exclusive amplification events on chr3 and ch4 across subclones.

We collect a large-scale 10x scDNA-seq dataset (10x_breast_S0) without known subclone labels, where 10,202 cells from five adjacent tumor dissections (A, B, C, D, and E) of triple-negative breast cancer are sequenced. We check whether SEAT seizes the substantial intra-tumor heterogeneity. In Fig. 4B-D, SEAT automatically detects seven subpopulations, and the proportions of the cell subpopulations vary across the five lesions. The blue subpopulation c6 gathers normal cells, with the mean cellular ploidy being diploid across all sections. The number of normal cells gradually decreases from sections A to E. SEAT identifies six clonal subpopulations (c0-c5), where c3 manifests the highest average ploidy. The mutually exclusive amplification events (marked with black boxes in Fig. 4D) on chr3 and chr4 of subclones c0, c1, c2, and c4, indicate an early branching evolution which is consistent with the findings of Wang *et al*. (68); that is, originated from normal cell group c6, the earliest subclone could be c5, with CN=3 on ch3 and ch4. Subclone c5 derived to subclone c0 with amplification on chr3 (CN=4). Moreover, subclone c5 derived to an intermediate subclone with amplification on chr4 (CN=4). Then, the intermediate subclone derived to subclone c1, c2, and c4 with CN gains on other chromosomes.

Furthermore, SEAT distinguishes cells with CNV gains and losses in circulating tumor cells of seven lung cancer patients (55) and in human cortical neurons (56) (Supplementary Fig. S28). SEAT also detects the loss of heterogeneity event, it successfully classifies chrX-bearing, chrY-bearing, and aneuploid sperm cells (57, 58) (Supplementary Fig. S28).

### Cell hierarchy dissects the chromatin accessibility heterogeneity of single-cell data

SEAT dissects chromatin accessibility heterogeneity of single cells. We utilize three public scATAC-seq data as benchmarking sets with gold standard cell type labels. scatac_6cl is a mixture of six cell lines (BJ, GM12878, H1-ESC, HL60, K562, and TF1) (59). Hematopoiesis consists of eight types of human hematopoiesis cells (CLP, CMP, GMP, HSC, LMPP, MEP, MPP, and pDC) (60). T-cell composes of four T-cell subtypes (Jurkat_T_cell, Naive_T_cell, Memory_T_cell, and Th17_T_cell) (61). We collect a multiome of scRNA and scATAC dataset, PBMC, for peripheral blood mononuclear cells (PBMCs) with 14 cell types.

The order of the cells in the agglomerative and divisive hierarchy is consistent with their ground truth cell types (Supplementary Fig. S29). The clustering accuracies of SEAT against its baselines are in Fig. 5A. In predefined-k mode, SEAT(k) demonstrates the highest clustering accuracies on scatac_6cl and T-cell sets. For auto-k clustering, SEAT(sub) beats Louvain and Leiden on all four sets. For scatac_6cl and T-cell, the optimal number of clusters obtained by SEAT matches the ground truth, thus yielding the comparable ARI against predefined-k clustering algorithms. Leiden and Louvain have lower performance due to predicting more clusters than ground truth (Supplementary Fig. S29).

**Fig. 5.**
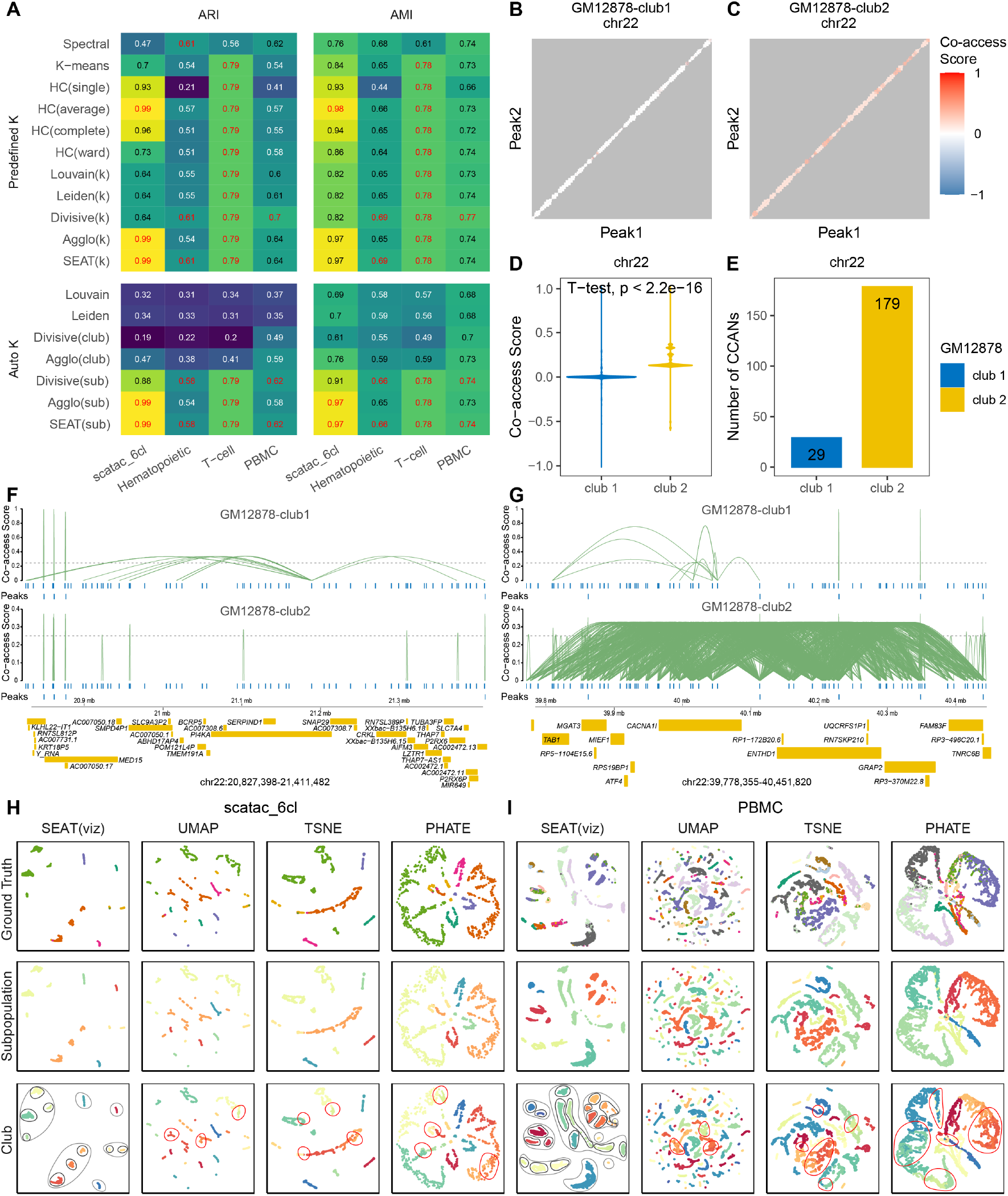
Applying SEAT on three scATAC datasets and one scRNA-scATAC multiome dataset. **A**. The adjusted rand index (ARI) and adjusted mutual information (AMI) of predefined-k and auto-k clustering tools. The best scores are colored red for each dataset in predefined and auto clustering benchmarking. Spectral: spectral clustering. HC(single), HC(average), HC(complete), and HC(ward): hierarchical clustering with single, average, complete, and ward linkage. Louvain(k) and Leiden(k): Louvain and Leiden in predefined-k mode. Divisive(k) and Agglo(k): the cell subpopulations from divisive and agglomerative hierarchy in predefined-k mode. SEAT(k): the cell subpopulations from SEAT cell hierarchy in predefined-k mode. Divisive(club) and Agglo(club): the cell clubs from the divisive and agglomerative hierarchy. Divisive(sub) and Agglo(sub): the cell subpopulations from divisive and agglomerative hierarchy in auto-k mode. SEAT(sub): the optimal subpopulations from SEAT cell hierarchy in auto-k mode. **B-D**. The co-accessibility score among peak pairs at chr22 for cells at SEAT club1 and club 2 from scatac_6cl GM12878 cell line. **E**. The number of *cis*-co-accessibility networks (CCANs) among pair of peaks at chr22 for cells at SEAT club1 and club 2 from scatac_6cl GM12878 cell line. **F**. The co-accessibility connections among *cis*-regulatory elements in chr22:20,827,398-21,441,482. The height of links signifies the degree of the co-accessibility correlation between the pair of peaks. The top panel illustrates cells in scatac_6cl GM12878-club1, and the bottom shows cells in scatac_6cl GM12878-club2. **G**. The co-accessibility connections among *cis*-regulatory elements in chr22:39,778,355-40,451,820. The height of links signifies the degree of the co-accessibility correlation between the pair of peaks. The top panel illustrates cells in scatac_6cl GM12878-club1, and the bottom shows cells in scatac_6cl GM12878-club2. **H**-**I** SEAT(viz), UMAP, TSNE, and PHATE plots of scatac_6cl and PBMC. The cells are colored with subpopulations, clubs, and ground truth. The gray and black circles in the SEAT(viz) plot indicate the subpopulation and club boundaries, respectively. In UMAP, TSNE, and PHATE plots, the red circles mark the unclearly segregated cell clubs. SEAT(viz): the hierarchical visualization from SEAT cell hierarchy.

We check whether SEAT reveals the functional diversity of single-cell chromatin accessibility. We select cells from scatac_6cl GM12878 cell line, then conduct *cis*-regulatory DNA interaction analysis on chr22 for SEAT cell club1 and club2. Fig. 5B-C depict the *cis*-regulatory map on chr22 of club1 and club2 cells, respectively. The co-accessibility correlations among peaks of club2 cells are significantly higher (*p* < 0.05) than club1 cells (Fig. 5D). Meanwhile, we identify 29 and 179 *cis*-co-accessibility networks (CCANs) from GM12878-club1 and GM12878-club2, respectively (Fig. 5E). The CCANs detected in GM12878-club1 and GM12878-club2 are heterogeneous. Fig. 5F illustrates a GM128780-club1 specified CCAN at chr22:20,827,398-21,441,482. The *cis*-regulatory elements surrounding gene *SNAP29* are co-accessible only in GM128780-club1. Moreover, we found dense pairwise connections among peaks at chr22:39,778,355-40,451,820 in GM12878-club2 (Fig. 5G), harboring genes *TAB1*, *MGAT3*, *MIEF1*, *CACNA1I*, *ENTHD1*, *GRAP2*, *FAM83F*, *TNRC6B*, *etc*.

Similar to the scRNA visualization refinement experiments, the SEAT(viz) reveals a clear pattern of cells corresponding to ground truth; and the nested layouts of subpopulations and clubs are clearly illustrated with gray and black circles (Fig. 5H-I and Supplementary Fig. S30). However, UMAP visualizations derived from high-dimensional data mix ground truth cell subpopulations in one clump (Supplementary Fig. S29). Furthermore, UMAP, TSNE, and PHATE visualizations derived from cell-cell similarity graphs fail to place cells from K562 (light green) and TF1 (yellow) within the vicinity in scatac_6cl; and they fail to place all effector CD8 T cells (magenta) together in PBMC (Fig. 5H-I). Likewise, the cell clubs marked with red circles are unclearly segregated in UMAP, TNSE, and PHATE plots.

## Discussion

Detecting and visualizing cellular functional diversity is essential in single-cell analysis. Neglection of the underlying cellular nested structures prevents the capture of full-scale cellular functional diversity. To address the challenge, we incorporate cell hierarchy to investigate the functional diversity of cellular systems at the subpopulation, club, and cell layers, hierarchically. The cell subpopulations and clubs catalog the functional diversity of cells in broad and fine resolution, respectively. In the cell layer, the order of cells further records the slight dynamics among cells locally. Accordingly, we establish SEAT to construct cell hierarchies utilizing structure entropy by diminishing the global uncertainty of cell-cell graphs. In addition, SEAT offers an interface to embed cells into low-dimensional space while preserving the global-subpopulation-club hierarchical layout in cell hierarchy.

Currently, state-of-the-art clustering tools for cell subpopulation or club investigation neglect the underlying nested structures of cells. Flatten clustering tools, such as spectral clustering (10) and K-means (11), do not support the cell hierarchy. Although conventional hierarchical clustering (12), Louvain (13) and Leiden (14) derive cell hierarchy layer by layer via optimizing merging or splitting metrics, computing these metrics merely uses single-layer information. When constructing subsequent layers, they have not incorporated the built-in cell hierarchy in the previous layers. Structure entropy is a metric that encompasses the previously constructed internal cell hierarchy. Experiments validate that SEAT delivers robust cell-type clustering results and forms insightful hierarchical structures of cells.

SEAT is good at finding the optimal subpopulation number with high accuracy. We have collected scRNA, scDNA, and scATAC profiles with the number of cell types ranging from 2 to 14. SEAT consistently predicts the optimal cluster number closest to the gold or silver standards, while Louvain and Leiden predict too many clusters. Especially for scRNA set Kumar, SEAT boosts the accuracy from 0.34 to 1 compared to Louvain and Leiden (Fig. 2A). Auto-k clustering mode of SEAT is comparable to or better than the best clustering results of predefined-k clustering methods for most datasets. SEAT specializes in hierarchically deciphering cellular functional diversity at subpopulation and club levels. We observe visible marker gene patterns that match cell clubs within one cell subpopulation. For the p3cl set, the basal, luminal, and fibroblast cell subpopulations have their own cell clubs, determined by differentially expressed cell cycle genes (*HIST1H4C*, *CDC20*, *CCNB1*, and *PTTG1*) (Fig. 2C). Looking at the seven agglomerative clubs for the basal subpopulation, we find a distinct breast cancer cell club that drives oncogenic *AREG*-*EGFR* signaling in all basal cells (Fig. 2D), suggesting a promoting role in tumorigenesis (67). Cell hierarchy obtained from copy number profiles of 10x_breast_S0 demonstrates a mutually exclusive sub-clones layout (Fig. 4D), indicating an early branch evolution (68). Furthermore, we find that there is a club-specified dense co-accessible network of *cis*-regulatory elements at chr22:39,778,355-40,451,820 in GM12878-club2, harboring genes *TAB1*, *MGAT3*, *MIEF1*, *CACNA1I*, *ENTHD1*, *GRAP2*, *FAM83F*, *TNRC6B*, *etc* (Fig. 5G).

Inferring the periodic pseudo-time for the cell cycle data is crucial as it reveals the functional diversity of cells undergoing the cell cycle process. Several tools are dedicated to cell cycle pseudo-time inference. CYCLOPS (15) and Cyclum (16) utilize deep autoencoders to project expression profiles into cell pseudo-time in the periodic process, which act as black boxes and lack explainability. reCAT (17) employs the Gaussian mixture model to group cells into clusters, and constructs a cluster-cluster graph weighted by the Euclidean distance between the mean expression profile of each cluster, then takes the traveling salesman path of the cluster-cluster graph as the order. Finding a traveling salesman path is NP-hard, and no polynomial time algorithms are available (17). CCPE (18) learns a discriminative helix to represent the periodic process and infer the pseudo-time. However, we fail to run CCPE according to its GitHub instruction. Moreover, CYCLOPS, Cyclum, reCAT, and CCPE bypass the nested structure of cells when inferring the pseudotime. In this study, we propose that the cell layer of a hierarchy encodes the pseudo-time of cells for cycling data. We build the hierarchy by minimizing the structure entropy of the kNN cell-cell graph. The built hierarchy carries the nested structure between individual cells and their ancestral cell partitions. Then, the order of individual cells is acquired with an in-order traversal of the hierarchy. scRNA data exemplify that SEAT cell orders outperform CYCLOPS, Cyclum, reCAT, and CCPE by accurately predicting the periodic pseudo-time of cells in the cell cycle process. In hESC and mESC cells, the expressions of M phase marker genes *UBE2C*, *TOP2A*, *CDK1*, and *CCNB1* rise progressively alongside the SEAT recovered order and are peaked at the M phase with significant fold changes (Fig. 3C).

Visualizing the hierarchical functional diversity of cells in biological systems is crucial for obtaining insightful biological hypotheses. UMAP (26) intends to maintain the global cell structures by minimizing the binary cross entropy. TSNE (27) preserves the local cell structures. PHATE (28) tackles the general shape and local transition of cells. However, none of them impart the nested structures of cells into the visualization. We propose a nonlinear dimension reduction refinement based on UMAP by incorporating cell hierarchy as supervised knowledge. We acquire three cell-cell graphs that only store the intra-connections of cells within each global, subpopulation, and club partition. Then, we minimize the weighted binary cross-entropy of the three cell-cell graphs. This approach guarantees the global structure of the cells. Moreover, it ensures that cells within one cell club and cell clubs within one subpopulation are closely placed in the visualization. In contrast, cells from different clubs and subpopulations are kept at a considerable distance. One can adjust the cross-entropy weights of global-subpopulation-cell layers so that the patterns in visualization retain a desired degree of hierarchy. Experiments with scRNA and scATAC data demonstrate that SEAT hierarchical visualization consistently produces a clear layout of cell clumps corresponding to the cell hierarchy.

Cellular abnormalities may distort the entire cell hierarchy. When there are cell outliers presented, the original SEAT will assign each cell outlier to its nearest cell subpopulation. Thus, the downstream biological interpretation may be skewed. To tackle the issue, we provide an optional average kNN outlier detection step before constructing the cell hierarchy. In Supplementary Results and Supplementary Fig. S31-S35, we demonstrate the distance cutoff is more stable than the distance percentile cutoff because the latter heavily depends on the ratio of outliers in the whole population. Thus, we set distance cutoff as the default outlier detection strategy.

The structure entropy evaluates the global uncertainty of random walks through a network with a nested structure (19). The minimum structure entropy interprets a stable nested structure in the network. Li *et al*. has used structure entropy to define tumor subtypes from bulk gene expression data (21) and to detect the hierarchical topologically associating domains from Hi-C data (22). These works utilize greedy merging and combining operations to build a local optimal multi-nary hierarchy and cutting hierarchy roughly by keeping the top layers. As we have proven that a binary hierarchy of minimum structure entropy exists for a graph (23), Li *et al*.’s strategy to search for a multi-nary hierarchy is not optimized. Adopted by Louvain and Leiden, modularity is a popular optimization metric to capture community structure in a single-cell network. Agglo(club) is analogous to Louvain’s if we switch the merging metric to modularity. Agglo(club) achieves better or comparable clustering performance against Louvain in most benchmark sets (Fig. 2A and Fig. 5A), suggesting the superiority of structure entropy over modularity in measuring the strength of hierarchically partitioning a network into subgroups. We have discussed the differences and advantages of SEAT against the existing structure entropy and modularity approaches at the algorithmic level in the Supplementary Method.

SEAT detects the cell hierarchy, assuming that the entropy codes nested structures of cells. There is no assurance that the resultant cell hierarchy will resemble accurate nested structures of cells. SEAT finds a pseudo cell hierarchy of cells. We show that the pseudo cell hierarchy showcases profound efficacy and biological insights in subpopulation detection, cell club investigation, and periodic pseudo-time inference for single-cell multiomics benchmarking datasets. In future work, we aim to refine the algorithm to find a more accurate and insightful pseudo cell hierarchy.

Recall that the cell hierarchy has multiple layers to present cellular heterogeneity. In this study, we merely utilize four main layers (global, subpopulation, club, and cell) to interpret and visualize the cellular functional diversity. In the future, we intend to investigate possible biological insights and visualization layouts derived from more cell hierarchy layers.

Moreover, the order of the cell clubs can be flipped in the cell hierarchy. There is only a partial order among cells bounded by the cell hierarchy. We plan to refine the algorithm to provide a proper non-partial one-dimensional order, which might infer the nuance of pseudo-time or development trajectory among cells outside the periodic cell cycle.

## Data Availability

The 25 scRNA, seven scDNA, three scATAC, and one scRNA-scATAC multiome datasets are publicly available. The details are summarized in Experiment Setting and Supplementary Methods.

## Code Availability

The source code of SEAT is available at https://github.com/deepomicslab/SEAT.

## Funding

This project is supported by CityU/UGC Research Matching Grant Scheme 9229012.

## Author Contribution

LXC conducted the project and wrote the manuscript. SCL supervised the project and revised the manuscript.

## Notes

### Competing Interest Statement

The authors have declared no competing interest.

### Summary of Updates

Add outlier detection, language and figure refinements.

